# Global Effects of a PD Risk-SNP at the Alpha-Synuclein Locus

**DOI:** 10.1101/2021.07.06.451330

**Authors:** Jordan Prahl, Steven E. Pierce, Edwin JC van der Schans, Gerhard A. Coetzee, Trevor Tyson

## Abstract

**SUMMARY:** One of the most significant Parkinson’s disease (PD) risk variants, rs356182, is located at the PD-associated locus near the alpha-synuclein encoding gene, *SNCA*. *SNCA*-proximal variants, including rs356182, are thought to function in PD via allele-specific regulatory effects on SNCA expression. However, this interpretation discounts the complex activity of genetic enhancers and possible nonconical effects of alpha-synuclein. Here we investigate a novel risk mechanism for rs356182. We use CRISPR-Cas9 in LUHMES cells, a model for dopaminergic neurons, to generate precise hemizygous lesions at rs356182. The PD-protective (A/-), PD-risk (G/-), and WT (A/G) strains are differentiated into dopaminergic neurons then compared transcriptionally and morphologically. We observe effects not typically ascribed to *SNCA*; hundreds of differentially expressed genes associated with neuronal differentiation and axonogenesis. Together, the data implicate a risk mechanism for rs356182 in which the risk-allele (G) is associated with abnormal neuronal differentiation. We speculate the disease-relevant effect originates as a diminished population of DA neurons leading to the predisposition for PD later in life.

## INTRODUCTION

With each consecutive genome-wide association study (GWAS), Parkinson’s disease (PD) is associated with an ever-increasing number of single-nucleotide polymorphisms (SNPs) [1, 2]. Because these associations are based on large epidemiological studies, descriptions of risk mechanisms require follow-up experiments. For most PD risk loci the identification of the causal SNP(s) has not been definitive, with few notable exceptions [3–6]. Most GWAS-identified risk loci are located in non-coding regions of the genome, further complicating this process [7]. One of the the top GWAS-identified PD risk-SNP, rs356182, is an example of a prominent SNP lacking a confirmed mechanism.

Rs356182 has a meta p-value of 1.85×10^−82^, making it the most significant association among non-coding PD risk-SNP [8]. The risk allele (G) is robustly represented in the population (frequency=40-50%) but also has one of the highest odds ratios (1.34) of any PD-associated SNP [8, 9]. This means the proportion of G alleles in the PD population compared to healthy controls is unusually high even compared to statistically significant risk signals. Enticingly, this SNP is within a genetic enhancer and located close to *SNCA*, the gene encoding alpha-synuclein, which is one of the first proteins linked to familial PD as well as neuropathologically present in Lewy bodies (Figure 1A) [7, 10, 11]. These data have led to the assumption that rs356182 confers risk for PD by directly and even exclusively regulating the expression of *SNCA*. However, this narrow view of the rs356182 mechanism neglects the potential for additional enhancer-promotor interactions, as well as secondary and tertiary effects of transcription factor (TF) misregulation.

**Figure 1:**
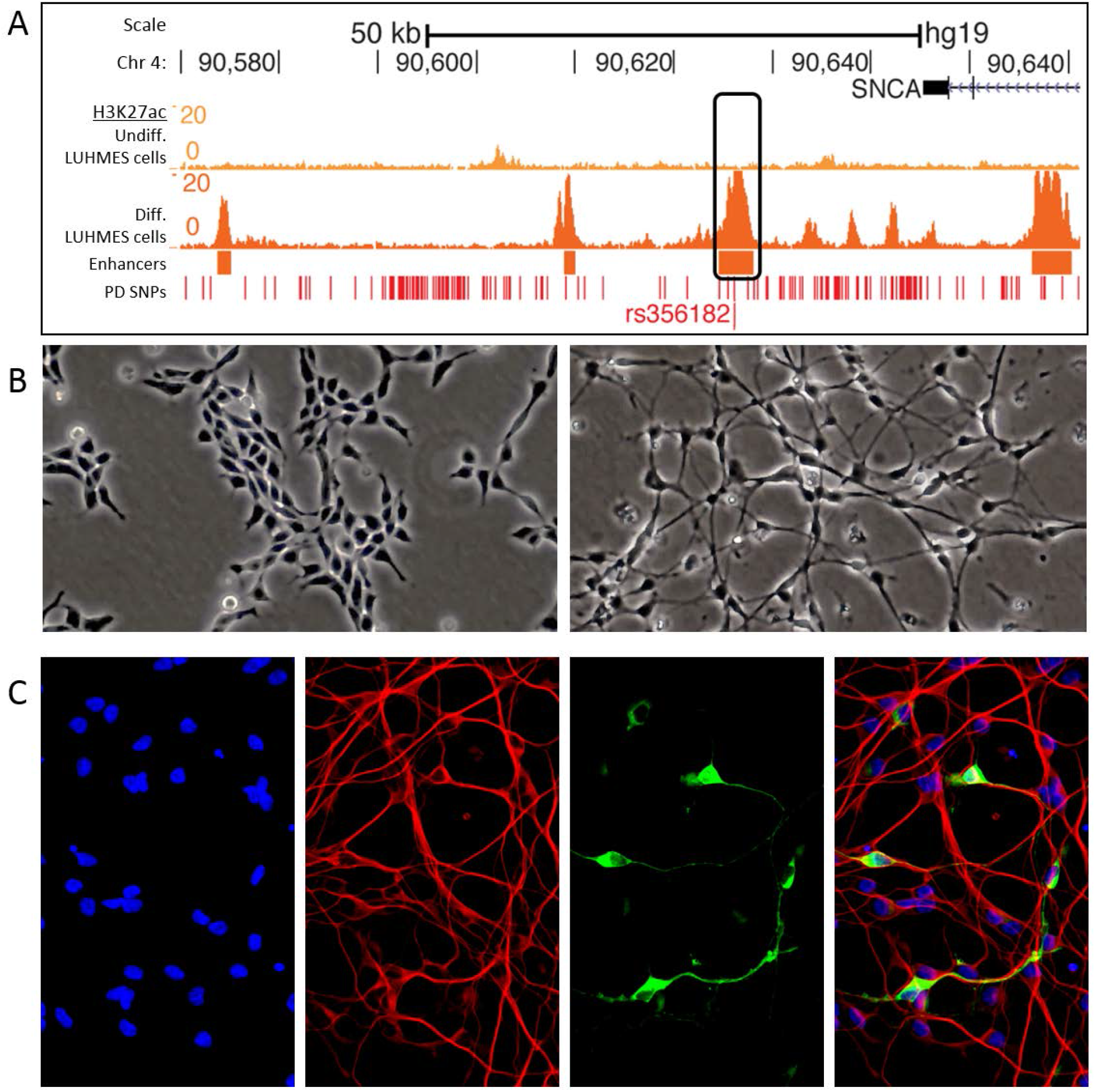
An enhancer encompassing rs356182 is active in differentiated LUHMES cells. (A) Histone H3K27ac track for undifferentiated (top) and differentiated (bottom) LUHMES cells. (B) Bright-field images showing the morphology of wildtype undifferentiated (left) and differentiated (right) LUHMES cells. (C) Immunofluorescent (TUJ1= red, and TH= green) and stained images (nuclei DAPI= blue) of differentiated LUHMES cells (day 6 of differentiation).

Unlike genes and promotors, enhancers are highly dynamic genetic elements, in the sense of functioning both cryptically via multiple TFs and with different gene targets in different cell types and stages. Enhancer variants have continuous effects (graded responses on gene expression) as apposed the binary effects of gene triplet code variation. Furthermore, enhancers can interact with the closest gene promotor on linear DNA (*SNCA* in this case). However, up to two-thirds of enhancers skip the nearest gene entirely and interact exclusively with distal promotors [12, 13]. Additionally, enhancers may interact with multiple promoters, and a single promoter often interacts with numerous enhancers in the three-dimensional space of chromatin [14, 15]. Enhancers regulate the expression of target genes by recruiting TFs to specific regulatory binding motifs. In this way, SNPs may modulate downstream gene expression by changing the binding affinity of TFs to a particular allele and alter the strength of the enhancer-promotor interactions.

Lund Human Mesencephalic (LUHMES; a.k.a. MESC2.10) cells offer a unique platform for studying immature human midbrain neurons and the epigenetic architecture present during neuronal development, due to their isolation from 8-week old fetal mesencephalic tissue [16]. LUHMES cells are immortalized in a stem-like state with the v-MYC oncogene, which, when deactivated allows the cells to enter the differentiation pathway and reach their terminal state within 6 days. In their differentiated state, LUHMES cells have a neuronal phenotype, displaying long neuronal projections (Figure 1B), and express the rate-limiting enzyme in dopamine production, tyrosine hydroxylase (TH) (Figure 1C). Beneficial to this research, LUHMES cells are heterozygous for rs356182 with one chromosomal homolog possessing the PD risk-conferring allele, guanine (G), and the other homolog possessing the protective allele, adenosine (A), on the Watson strand. Interestingly, when differentiated, these cells also contain an enhancer surrounding the rs356182 SNP [7]. This makes LUHMES cells an ideal model for investigating PD pathology or predisposition originating very early in development, including neuronal differentiation-based mechanisms.

This study seeks to elucidate the mechanism surrounding rs356182 by examining allele-specific gene regulation and changes to differentiated morphology associated with this SNP. To that end, we used CRISPR-Cas9 to generate mono-allelic lesions of rs356182 in LUHMES cells (Figure 2A), and examined the changes in gene expression and morphology therein. We observed a novel rs356182 risk mechanism not previously attributed to *SNCA*, pertaining to neuronal differentiation processes.

**Figure 2:**
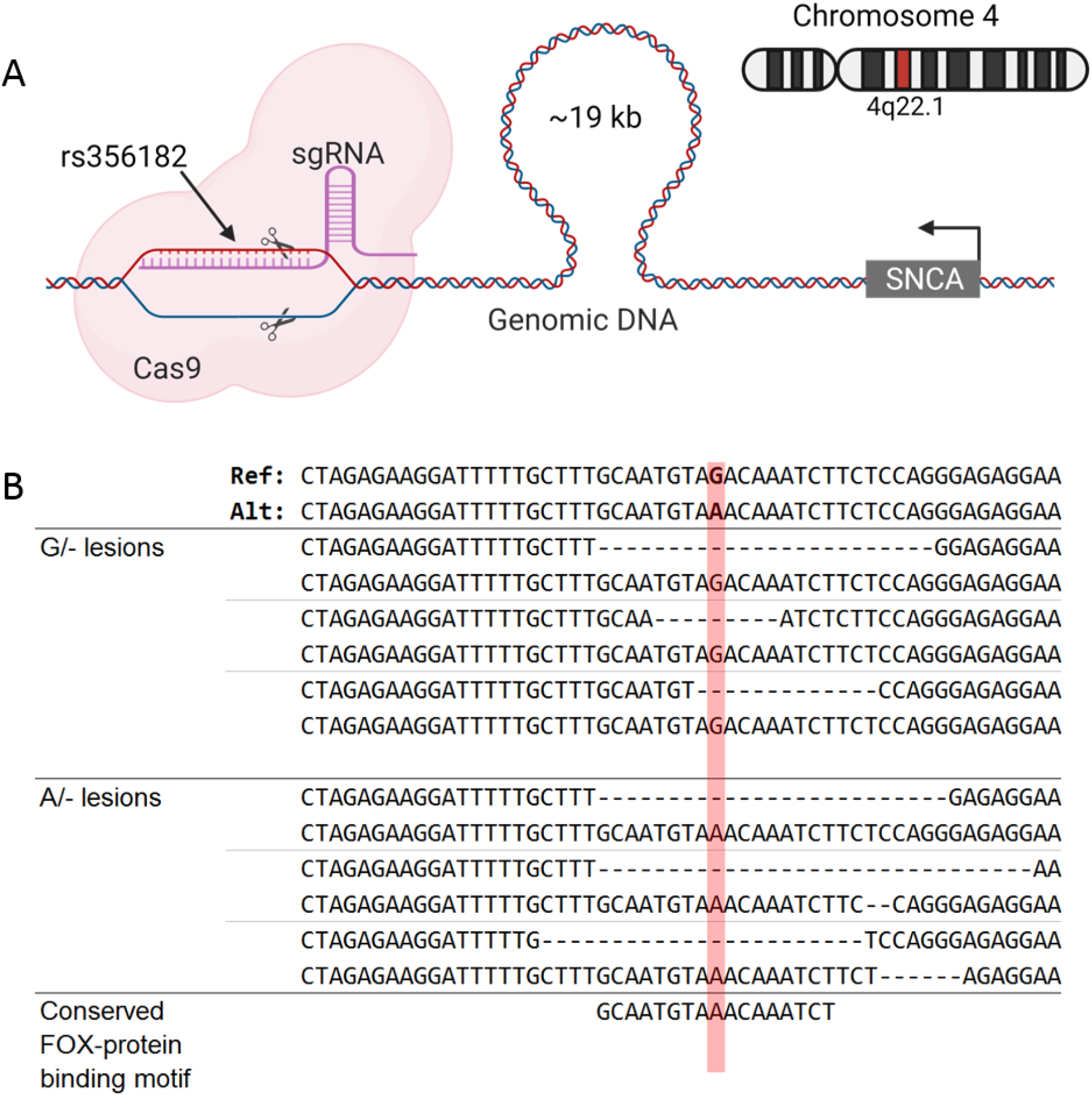
Excised rs356182 by CRISPR-Cas9. (A) Schematic depicting the position of rs356182 relative to *SNCA* on chromosome 4 and the guide-RNA targeting that locus for a double stranded DNA break. (B) The resulting deletions (deleted bases depicted as dashes) from targeting rs356182 with CRISPR-Cas9; position of rs356182 highlighted in red.

## METHODS

### LUHMES cell model

LUHMES cells, obtained from ATCC (CRL-2927), were cultured essentially as described by Scholz et al. [17]. As previously described [7], the cells were incubated in a humidified 37°C, 5% CO2 incubator in flasks pre-coated with 50 mg/mL poly-L-ornithine (Sigma, Cat # P3655) and 1 mg/mL fibronectin (Sigma, Cat # F114) in water. The coated flasks were incubated at 37°C overnight, rinsed with water, and allowed to dry before seeding cells. Cells were cultured in complete growth medium containing Advanced DMEM:F12 (Thermo Fisher, Cat # 12634-010) with 2mM L-glutamine (Thermo Fisher, Cat # 25030081), 1X N-2 supplement (Thermo Fisher, Cat # 17502-048), and 0.04 mg/mL bFGF (Stemgent, Cat # 03-0002). Cells were allowed to reach 80% confluency before passaging with 0.025% trypsin/EDTA. Before differentiation, cells were seeded at 3.5×10^6^ per T75 flask containing complete growth medium and incubated at 37 °C for 24 h (day -1). For induction of differentiation, culture medium was changed to freshly prepared DMEM:F12 with 2mM L-glutamine, 1X N-2, 1mM cAMP (Carbosynth, Cat # ND07996), 1 mg/mL tetracycline (Sigma, Cat # T7660), and 2 ng/mL glial cell line-derived neurotrophic factor (GDNF) (Sigma, Cat # G1777) (day 0). LUHMES cells then grow in the differentiation media for 2 days before being passaged, again into differentiation media (day 2). Finally, cells were harvested on day 6 after the initial introduction of differentiation media (day 6).

### CRISPR-Cas9 Editing

For each target sequence, double-stranded DNA sequences complementary to the target sequences were generated by PCR and then cloned into the pSpCas9(BB)-2A-GFP (PX458) vector (AddGene, cat. # 48138) (Table S1). Ligated plasmids were delivered into Stbl3 chemically competent E. coli cells and selected from ampicillin-treated auger plates. Transformed E. coli colonies were isolated and expanded in LB media. Expanded plasmids were purified using the QIAprep Spin MiniPrep Kit (Qiagen, cat. # 27106). LUHMES cells were electroporated and transfected with 2 μg of plasmid using the Amaxa™ Basic Primary Neurons Nucleofector™ Kit and protocol (Lonza, cat. # VPI-1003). Transfected cells were selected by flow-sorting for DAPI negative, GFP positive singlets into pre-coated 96-well plates (Figure S1). Sorted cells were clonally expanded and screened via Sanger DNA sequencing from Genewiz. Individual allele sequencing was achieved by TOPO-TA cloning (Thermo Fisher, cat. # K4575J10), followed again by Sanger sequencing. Clones with a confirmed edit disrupting rs356182 were used in the edited conditions, while sorted clones which did not have a confirmed edit at rs356182 were used as the A/G lesion controls (Figure 2B).

### RNAseq

Three clones of each of the G/- lesion model and A/- lesion model and parent (wild-type) G/A clones were selected for RNAseq analysis (Table S2). The three wildtype control clones (described in the CRISPR-Cas9 editing protocol above) were also selected for RNAseq, as well as 9 true wildtype clones (unexposed to CRISPR and dilution sorted) in both their undifferentiated and differentiated states. Clones were differentiated following standard LUHMES cell differentiation protocol. Clonal RNA was isolated and purified using the QIAGEN QIAshredder (Cat. #: 79654) and RNeasy isolation kit (Cat. #: 74104). Total RNA was submitted to the Van Andel Research Institute’s Genomics Core for QC, library preparation, and paired-end sequencing using the Illumina NovaSeq 6000 in split-lane SP Flowcells. Sequencing results were saved as zipped FASTQ files.

### Differential gene expression

First, Illumina sequencing adapters were trimmed from the raw count files using the TrimGalore software package (version-0.6.0). Next, trimmed reads were aligned to the GRCh37-hg19 reference genome using the STAR software (Spliced Transcripts Aligned to a Reference; version- 2.7.8a). After the alignment, sub-threshold count reads were filtered out and raw counts were normalized to count per million (CPM) and trimmed mean of M values (TMM normalization) using EdgeR (version-3.32.1). Samples were submitted with a minimum biological triplicate; therefore, genes expressed in fewer than 3 samples were excluded. A z-score was determined for each gene per sample and fold-change and adjusted P-values were calculated using the limma-voom method (version- limma-3.46.0. Detailed scripts on Github.

### Gene Set Enrichment Analysis (GSEA)

GSEA for biological processes was performed using ClusterProfiler (version- 3.18.1). The background list of genes in the universe was described as the full list of genes expressed in our LUHMES models. In each case, an adjusted P-value < 0.05 was the maximum threshold for modulated genes. Gene sets were separated by condition and increased/decreased gene expression. An absolute fold-change greater than 2 was used as the minimum threshold for modulated genes. To reduce redundancy of ontological terms and the bias introduced by over-represented genes, terms that have greater than 70% of shared gene annotations were collapsed into one group.

### Morphology

For morphological analysis, clones were differentiated following the standard LUHMES cell differentiation protocol. On day-2 of differentiation, clones were passaged to the standard density of 1.5×10^5^ cells/cm^2^ into pre-coated 24-well plates. On day 6 of differentiation, clones are fixed by incubating in 2% formaldehyde solution for 5 minutes, followed by incubation in 4% formaldehyde solution for 15 minutes, then stored in DPBS at 4°C for up to 2 weeks. Immunofluorescence was conducted as described by the Invitrogen Human Dopaminergic Neuron Immunocytochemistry Kit (Catalog no. A29515). In brief, cells were incubated in perm/block buffer (1% BSA, 0.3% Triton X-100, and DPBS) for 30 minutes at room temp. Primary antibody was added directly to the perm/block buffer in the wells and incubated overnight at 4°C. Wells were washed 3x in DPBS for 2 minutes at RT, and then secondary Ab was added to the cells for 1h at RT (Table S3). Wells again were washed 3x in DPBS. On the third wash, 1-2 drops/ml of DAPI was added to the wash buffer. Cells are then stored in DPBS for <2 weeks.

After immunohistochemical and DAPI staining, samples were imaged on the Zeiss Celldiscover7 (CD7) at 10x magnification (20 Z-stacks) by the VARI Optical Imaging Core. Composite images were prepared using the ZEN 3.2 (blue edition) software by using automated threshold settings for the DAPI and TUJ1 channels and manually setting the TH channel and z-stack. The TH channel must be manually set due to the skewed intensity distribution disrupting the automated threshold settings. Split fluorescent channels were converted to binary images in ImageJ and analyzed for pixel coverage and cell count. Measurement of cell density was calculated as the percent of the well covered by DAPI stain (binary images; Black pixels / total pixels). Neurite development was calculated as the ratio of cell body stain (TUJ1) and nuclei stain (DAPI) (binary images: black pixels / total pixels, Red channel / blue channel).

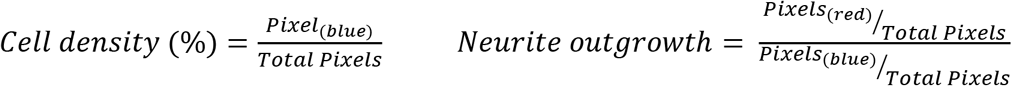

## RESULTS

### Disruption of rs356182 has global effects and allele-specific activity

We sought to determine if rs356182 is indeed functional and identify the potential allele-specific activity of this SNP. To that end, we created hemizygous clones at this site and therefore possessing a lesion (i.e. short deletion) on one chromosome while maintaining the original major (G) or minor (A) alleles on the other. To achieve this, we used CRISPR-Cas9 to target the PAM site nearest to rs356182 (Figure 2A & Table S1). In individual cells, lesions were randomly formed at either the A-allele or G-allele and were approximately 20 ± 8 bp following DNA repair (Figure 2B). After screening, 3 clones for each of 3 conditions were generated; the PD-protective condition which maintains only the protective allele (A/-), the PD-risk condition which maintains only the risk allele (G/-), or the wildtype control condition in which clones show no edits and maintain both rs356182 alleles (A/G). Statistical analysis comparing the length of lesions replicates in the protective and risk conditions indicated no significant difference in the average deletion size (mean leasion 20 bp, G/- vs A/-, Student’s T-test: P=0.1176, Wilcox test: P=0.20). Clonally derived hemizygous (single allele) strains and wildtype (both alleles) controls were differentiated and allele-specific gene expression was examined using RNA-seq. As the deletion method provided slightly different-sized lesions, expression results were analyzed for relation to lesion size but showed no correlation. We found many and widespread differentially expressed genes, contrary to the expectation that *SNCA* would be the primary gene target. Whereas *SNCA* was significantly modulated in both hemizygous conditions compared to WT (Figure 3A), neither *SNCA* nor any other genes near rs356182 were the most significant or highest fold-changed genes affected (Figure 3B). Our results here corroborate previous results which showed a higher *SNCA* expression in the A/A genotype and lowest expression in the G/G genotype [18]. We observed allele-specific differential gene expression spanning the entire genome (Figure 3C). We used principle component analysis (PCA) of gene expression for the 9 samples in order to visualize the systematic variance in transcription. The results show that the protective, A/- and A/G conditions were closely space along principle component 1 (accounting for 34.4% of the sample variance), and imply that the PD-protective condition is more similar to the wild-type condition than it is to the risk condition (Figure 3D). Ultimately, both conditions showed several hundred genes significantly altered compared to the wildtype controls, including both up and down-regulated genes with absolute fold-change greater than two (Table S5).

**Figure 3:**
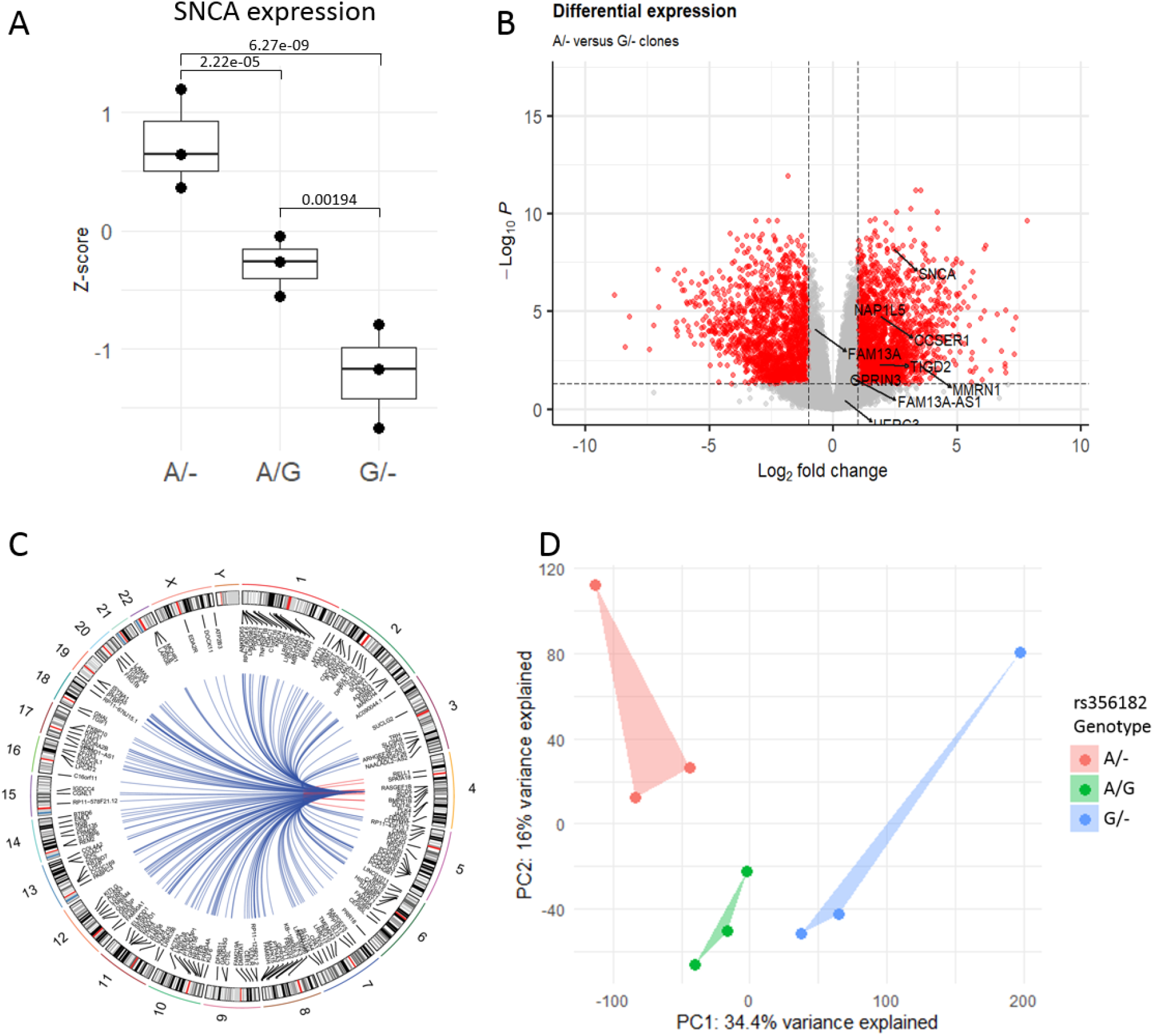
Disruption of rs356182 has global effects and allele-specific activity. (A) Box-and-whicker plot of z-scores for SNCA gene expression; adjusted P-value displayed above each group comparison. (B) Volcano plot showing the differentially expressed genes between the A/- and G/- clones with the genes proximal to rs356182 annotated; Log_2_ fold change cutoff, 1; p-value cutoff, 0.05. (C) Circos plot depicting the significantly modulated genes between the A/- and G/- clones across the entire genome. (D) PCA plot showing principle components 1 (X-axis) and 2 (Y-axis).

### CRISPR-mediated lesions at rs356182 modulate the capacity for differentiation

Based on the specific activation of the rs356182-containing enhancer during the differentiation of DA neurons (Figure 1A) [7], we hypothesized that the elements at this locus are involved in the differentiation process. Neuronal differentiation is a complex biological process and determines cell proliferation, morphology, metabolism, and communication (among many other processes). We have previously shown that the differentiation of LUHMES cells has a profound impact on gene expression, with thousands of genes significantly modulated between states of differentiation [7]. As expected, gene set enrichment analysis (GSEA) of those impacted genes showed enrichment in GO terms associated with neuronal differentiation and neurogenesis (upregulated genes), and cell cycle (down-regulated genes) [7]. Relative to the extreme comparison of neuronal vs proliferating LUHMES, the lesions strains we generated here had more modest effects on differentiated gene expression (Figure 4 & Table S5). Unexpectedly though, our lesion conditionss also resulted in differential expression for gene sets enriched for several of the same neuronal developmental ontological branches (Figure 4 & Table S6). Specifically, genes upregulated in the protective condition (A/-) participate in the positive regulation of neurogenesis, axonogenesis, and neuronal differentiation. Meanwhile, genes down-regulated in the risk condition (G/-) are enriched in terms associated with axonogenesis, neuron projection guidance, and synaptic function. The conclusion is that differentiation and neuronal morphogenesis are enhanced in the protective condition and inhibited in the PD risk condition.

**Figure 4:**
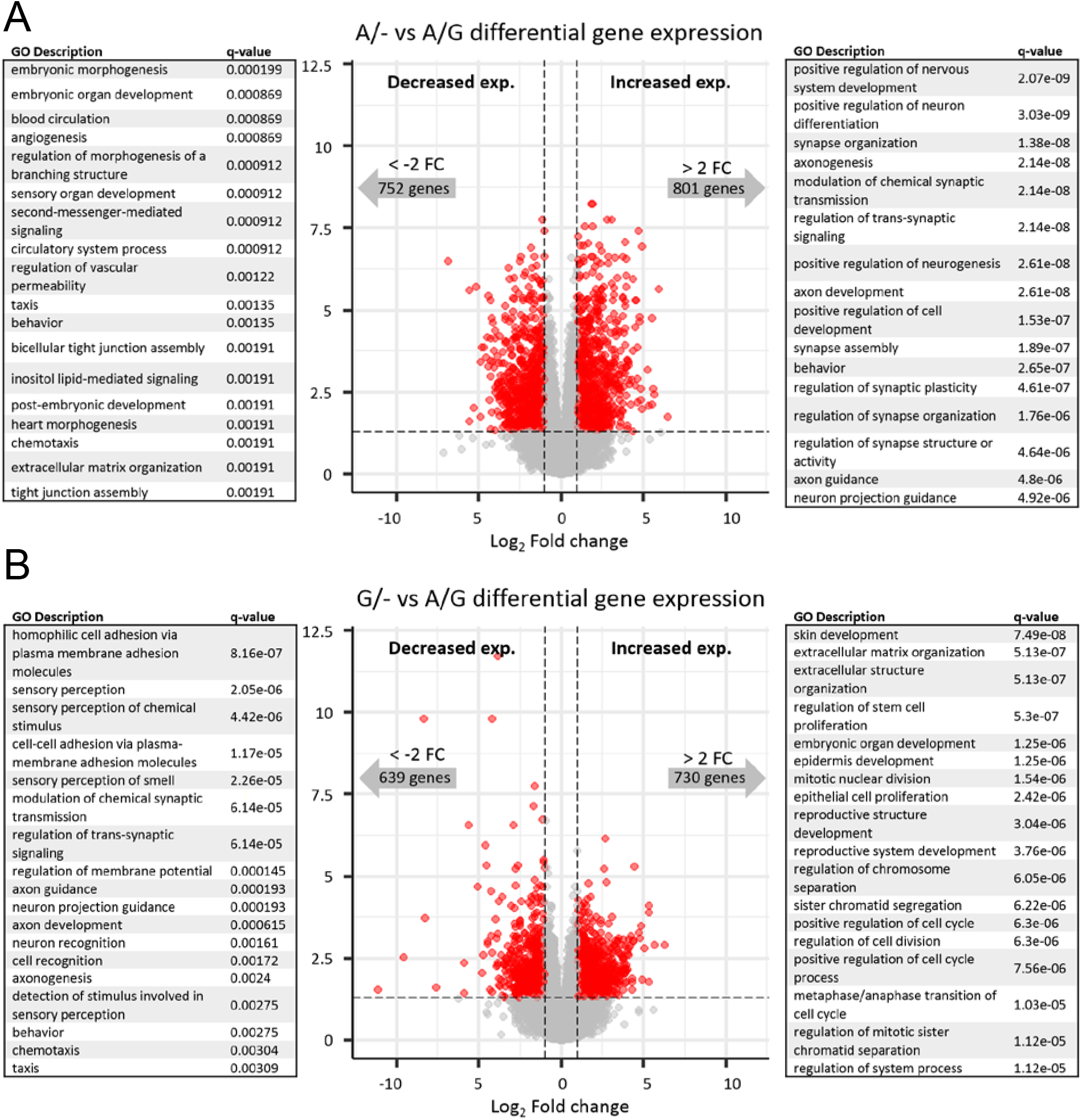
CRISPR-mediated lesions at rs356182 modulate the capacity for differentiation. Volcano plot of RNA-seq data depicting the differentially expressed genes between the lesion clones and wildtype clones. The grey dots represent genes that fail to meet minimum thresholds for adjust p-value (<0.05) and/or log_2_ fold change (>1). To the left and right, the top GO terms and associated P-values for each differentially expressed (up or down) gene set. (A) A/- versus A/G. (B) G/- versus A/G.

### The rs356182 risk-allele mediates a pathological morphology change

Following GSEA, it was clear that the neurodevelopment was generally altered in our lesion conditions. To determine whether those genetic changes resulted in obvious morphological differences or other differences in cell growth we differentiated and stained cells with DAPI, as a nuclear marker, and treated with anti-TUJ1 (neuron-specific class III beta-tubulin) which marks neuronal soma and neurites, and anti-TH (tyrosine hydroxylase) as an indicator of dopamine production. The G/- clones had significantly more cells per well on day 6 of differentiation than their wildtype counterparts (seeded at equal densities on day 2 of LUHMES cell differentiation) (Figure 5A & 5B). We believe this is due to a differentiation-specific modulation of proliferative activity because edited clones did not show different growth rates in their undifferentiated state (Figure S2). Additionally, the cell density within each well was more variable in the risk clone samples than in the protective clones or wild-type clones (Figure 5C). These wells more commonly had clusters of densely packed cells and then sections of sparsely populated cells (Figure 5A, middle). The TUJ1/DAPI stain ratio is also smaller in the risk (G/-) condition indicating reduced neurite growth per cell, in line with results from GSEA (Figure 5D). Finally, the protective clones had significantly more TH-expressing cells on day 6 of differentiation than either the wildtype controls or the risk clones (Figure 5E).

**Figure 5:**
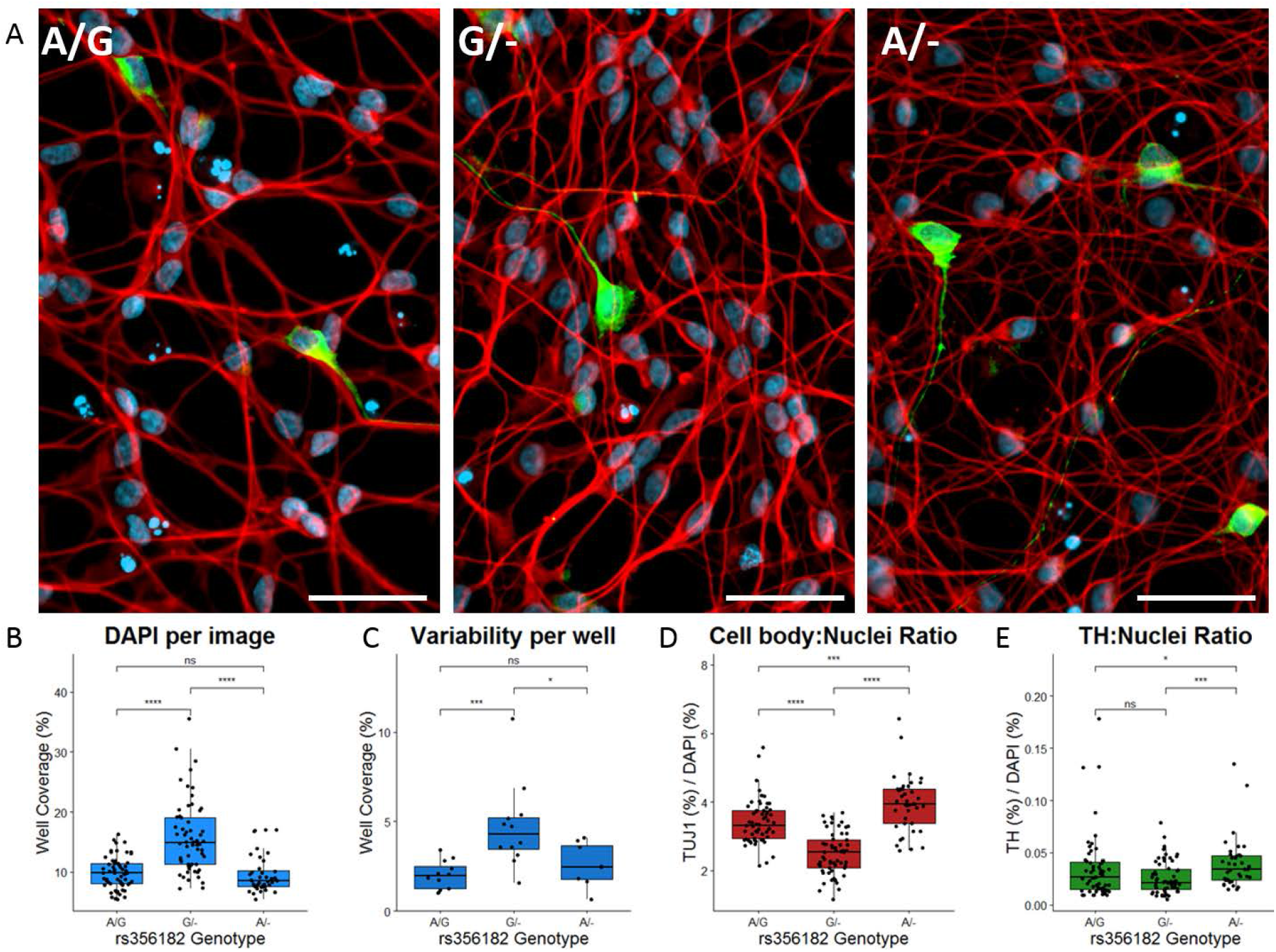
The rs356182 risk-allele mediates a pathological morphology change. (A) Representative immunofluorescent images from wildtype A/G clones (left), risk G/- clones (middle), and protective A/- clones (right); white bar= 50 μm, blue= DAPI, red= TUJ1, green= TH. (B-E) Quantification of immunofluorescent images by pixel coverage represented in box-and-whisker plots; Wilcoxon comparisons of nonparametric groups, ns: p>0.05, *: p<=0.05, **: p<=0.01. ***: p<=0.001, ****: p<=0.0001. (B) DAPI coverage per image as a measure of cell density. (C) Standard deviation of inter-well DAPI coverage. (D) Ratio of TUJ1/DAPI as a relative measure of cell body to nucleus. (E) Ratio of TH/DAPI as a normalized measure of TH-expressing cells within the cell population.

### rs356182 modulates transcription factor binding affinity

In situ analysis of the TF binding-motifs at the rs356182 region using HaploReg (Table S3) and MotifbreakR (Table S4) revealed a preferential binding affinity for the protective A-allele over the risk G-allele for the strongest interacting TFs at this locus (Tables S3 & S4) [19–21]. Notably, the Fox family of proteins was identified as strong candidate binders to the A-containing motif (Figure 2B & 6). Fox proteins regulate the expression of genes involved in branching morphogenesis (FOXA1/2), brain development (FOXA2/C1), and axon guidance (FOXD1), among a host of other biological processes [22]. The FOXA1 motif has previously been shown to be particularly vulnerable to variants that disrupt the 3 consecutive adenines, as the MotifbreakR results would suggest, and which is true of rs356182 (Figure 6) [23]. Additionally, the FOX family of proteins are known to engage in pioneer factor activity [24]. This data suggest that the rs356182 risk-allele (G) disrupts a particularly vital position within the putative binding motif, resulting in diminished binding affinity for FOX TFs, with the potential to reduce enhancer activity entirely.

**Figure 6:**
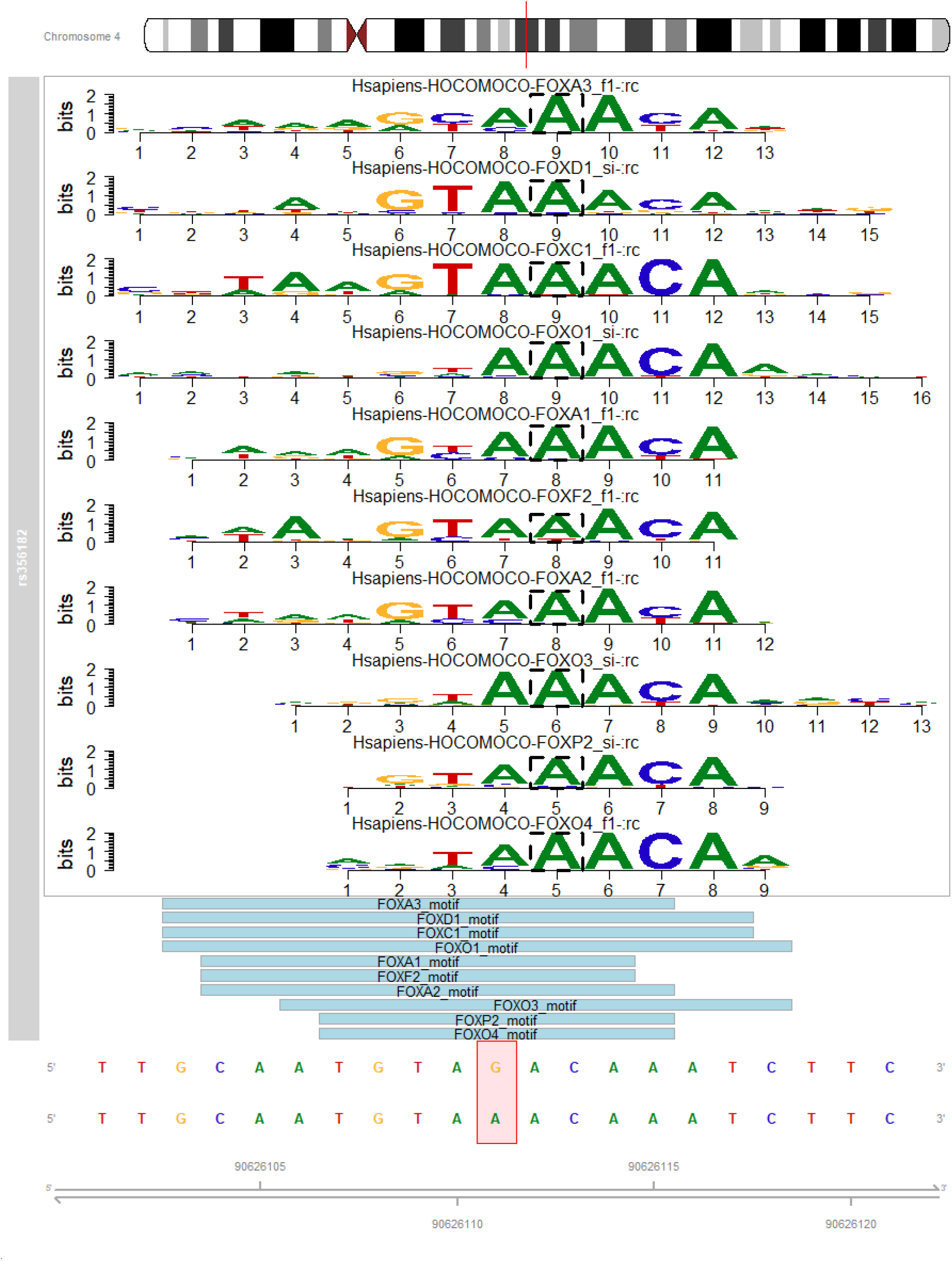
rs356182 modulates transcription factor binding affinity. (A) Results from MotifBreakR depicting the sequence and importance (letter height) of each base within the motif sequence for the “Strong interaction” proteins at this locus.

## DISCUSSION

There are more than a million genetic variants in the human genome, occurring on average once every six base-pairs depending on the minor allele frequency in the population. These variants, the majority of which are SNPs, have been used in GWASs to identify loci associated with a variety of traits and diseases, including PD. The value of the results obtained from these associations has been widely debated [25, 26]. The reason for this debate is that the mechanisms underpinning the associations are not always clear and most (>90%) of the risk SNPs don’t reside in protein coding regions, but instead in presumptive regulatory elements, which affect the expression of often unknown genes. Furthermore, many SNPs at a given locus are in linkage disequilibrium with each other, making causality assignment unclear. As of September 2018, the NHGRI-EBI GWAS catalog contains 5,687 GWASs of 71,673 variant-trait associations [27]. The vast majority of them provide a landscape view of possible functional involvements, leaving the majority with unknown functional significance. A major gap in our ability to understand and utilize GWAS results is thus the lack of in-depth mechanistic insights.

A rare example of mechanistic detailing of risk variant/gene association involves the variant within an intron of *FTO* and associated with increased risk for obesity and type-2 diabetes (T2D) [28]. Since epidemiological associations and data from mouse models indicate that *FTO* expression levels influence body mass, it seemed likely that the *FTO* intronic variants impose T2D risk via *FTO* expression regulation affecting enhancer activity. However, contrary to expectations, it was shown that that the *FTO* intronic variants are functionally and structurally connected with the homeobox gene *IRX3*, some mega-bases distant on linear DNA. It was demonstrated that IRX3 has a roll in controlling body mass composition and metabolism and represents a novel therapeutic target for diabetes.

Here we present evidence regarding the mechanism(s) surrounding rs356182 that contradicts the assumption that *SNCA* is the only affected gene. This assumption is that rs356182 directly and exclusively regulates the expression of *SNCA*, and in this way confers risk for PD. We show by the mono-allelic deletion of rs356182 that hundreds of genes, spanning the entire genome, are affected by rs356182. However, it remains unclear if these downstream gene changes are primary, secondary, or tertiary events. Furthermore, these affected genes appear to be enriched in GO terms related to the differentiation-proliferation processes. Our results suggest that: (i) *SNCA* has an important role as a regulator of DA neuron differentiation and that by manipulating rs356182, a cascade of genes is impinged by the resulting *SNCA* misregulation, or (ii) rs356182 does not exclusively regulate *SNCA* and is independently relevant to DA neuronal differentiation via the regulation of many genes. One, or both, of these scenarios, is likely true of the mechanism underlying the PD risk associated with rs356182.

To function as a transcription factor, α-SYN would need to be present at the site of transcription-- the nucleus. α-SYN was observed in both the nucleus and cytosol of neurons as early as the 1980’s, hence the name, but the implications of this localization have since been debated [29–31]. Under pathological conditions, such as in A53T *SNCA*-mutant containing neurons, nuclear localization of α-SYN is increased [32–34]. The α-SYN translocation is mediated by retinoic acid [35], and the accumulation and neurotoxicity are regulated by TRIM28 [36]. Once inside the nucleus, the mode of α-SYN associated neurotoxicity is less clear. α-SYN has been reported to bind to histones and increase fibrillation [32], bind to DNA directly to mediate DNA repair [37, 38], and interact with retinoic-acid response elements to regulate associated gene transcription which is linked to PD [35]. The DNA-repair mechanism associated with α-SYN has been debated and may be a result of normal cell response to DNA damage induced by α-SYN toxicity [38–40]. While it is clear that nuclear localization is linked to neurotoxicity [33], evidence for *SNCA* behaving as a TF is speculative.

The results from the motif analysis of rs356182 (Figure 1C, Table S3, & Table S4) imply that the A-allele is the primary functional allele and that the FOX family of proteins are the primary TFs binding to the A-containing motif. rs356182 is in the middle of the strong triple-A motif, which has previously been shown to be particularly important for TF binding, disruption of which having pathological consequences [23, 41]. Since FOX proteins have pioneer TF abilities, they are likely partially or wholly responsible for activation of the encompassing enhancer, resulting in the regulation of many genes by both primary- and secondary−/tertiary-interactions. What remains less clear is what is happening in the presence of the G-allele.

Based on the mechanism proposed above, deletion of the G-allele should have few, or no, consequences on gene expression, but this is not what we observed. In fact, deletion of the G-allele in the PD-protective clones promoted neuronal differentiation and neurogenesis (Figure 3). The fact that losing one allele increases the expression of certain genes and processes indicates that, contrary to what we predicted from binding-motif analysis, the G-allele is not simply a less functional allele, but is counteractively working against the A-allele. Further evidence that this is not a simple dose-response is that ~21% of the nearly 25,000 genes in our dataset have opposite expression changes between experimental conditions (i.e., one edited condition increases while the other edited condition decreases in expression compared to the wildtype), an example of which is *SNCA* itself (Figure 2A). Additionally, morphological analysis of body-nucleus ratio and the ratio of TH-expressing cells per condition showed the A/- clones seemingly differentiating more effectively than the risk clones, and even the wild-type A/G clones (i.e., A/- clones grew more/larger neurites per cell and more cells reached dopaminergic maturity within the timeframe). In other words, the A- and G-alleles impinge on the differentiation-proliferation mechanism by actively working against each other; the A-allele pushing the cell towards differentiation, and the G-allele holding it back in a more stem-like state. The dual functionality of the rs356182 alleles could explain why this locus has such a strong association with PD.

The PD field has known about the significant association of rs356182 to PD since the earliest GWASs due to its strong association with risk. Until now, the mechanism surrounding this risk was attributed to allele-specific expression of *SNCA*, variation of which would presumably predispose individuals to PD in their later years. However, we have identified a novel mechanism in which rs356182 impinges on the differentiation/proliferation mechanism during development. These results contribute to the growing body of evidence that PD is, at least in part, a developmental disorder [42, 43]. We speculate that rs356182 regulates neuronal differentiation and a cascade of related processes, leading to a diminished population of healthy dopaminergic (DA) neurons in individuals with the risk allele, making the individual more sensitive to subsequent insults to the DA cell population, and in this way confers risk for PD later in life [44].

## ACKNOWLEDGEMENTS

This research was supported in part by the Van Andel Research Institute (Grand Rapids, MI) Bioinformatics and Biostatistics Core, Flow Cytometry Core, Genomics Core, and Optical Imaging Core.

## AUTHOR CONTRIBUTIONS

**CRediT Taxonomy:** Conceptualization, JP, SEP, GAC, and TT; Methodology, JP and TT; Software, JP and SEP; Formal Analysis, JP and SEP; Investigation, JP; Resources, JP, EJCV, and TT; Writing-original draft, JP; Writing-Review & editing, JP, TT, SEP, and GAC; Visualization, JP and SEP; Supervision, GAC and TT; Funding Acquisition, GAC.

