## Supplemental Figures for "Global Effects of a PD Risk-SNP at the Alpha-Synuclein Locus"

**Table S1:** CRISPR Guides. rs356182 denoted in red.

| *Guide ID* | *Guide Sequence* | *Location (Hg19)* | *Model* |
|---|---|---|---|
| rs356182_A-allele | AATGTAAACAAATCTTCTCC | Chr4: 90625991 | Lesion (G/-) |
| rs356182_G-allele | AATGTAGACAAATCTTCTCC | Chr4: 90625991 | Lesion (A/-) |

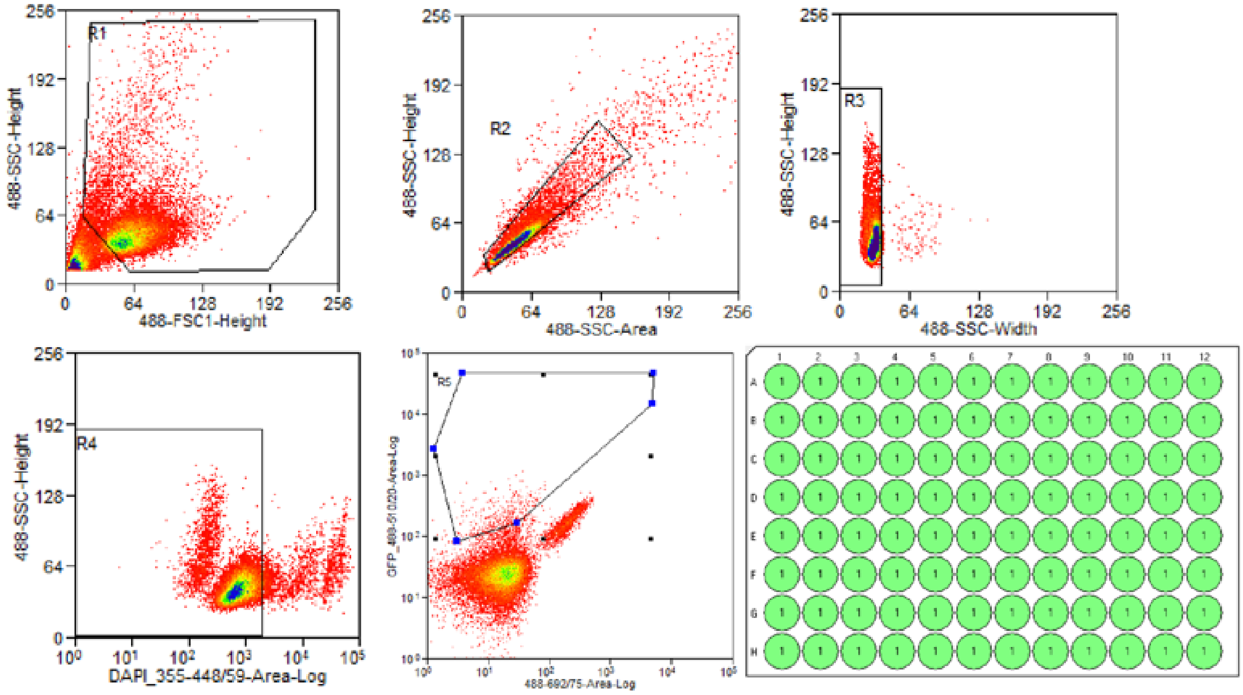


**Figure S1:** Flow sorting schema. From top, left-to-right, filter out cell debris, singlet selection, further singlet selection, live cells (DAPI negative), GFP positive (plasmid transfected), and plate setup.

**Table S2:** Antibodies and dilutions for morphological imaging.

| *Antibody* | *Primary or secondary* | *Staining dilution* | *Cat. #* | *Source* |
|---|---|---|---|---|
| Ab Rabbit Tyrosine Hydroxylase | 1 | 1:250 | P40101-150 | Pel-Freez |
| Purified anti-Tubulin β 3 (TUBB3) Antibody | 1 | 1:1000 | 801201 | BioLegend |

| | | | | |
|---|---|---|---|---|
| Goat anti-Mouse IgG (H+L) Highly Cross-Adsorbed Secondary Antibody, Alexa Fluor 594 | 2 | 1:250 | A11032 | Invitrogen |
| Goat anti-Rabbit IgG (H+L) Cross-Adsorbed Secondary Antibody, Alexa Fluor 488 | 2 | 1:250 | A11008 | Invitrogen |

**Table S3:** Full list of transcription factors with significantly modulated binding affinities at the rs356182 locus from Haploreg (version-4.1). (Library from Kheradpour and Kellis, 2013)

| *Position Weight Matrix ID* | *Strand* | *Ref* | *Alt* | ***Match on:***<br>Ref: CTAGAGAAGGATTTTTGCTTTGCAATGTA**G**ACAAATCTTCTCCAGGGAGA<br>Alt: CTAGAGAAGGATTTTTGCTTTGCAATGTA**A**ACAAATCTTCTCCAGGGAGA |
|---|---|---|---|---|
| *AP-3* | - | 10.8 | -1.1 | WRTKKRGR |
| *Fox* | - | 9.6 | 13.1 | WAARYAAAYAWWV |
| *Foxa_disc1* | + | 3.6 | 15.4 | WRWGYAAAYA |
| *Foxa_known1* | - | 4.5 | 11.8 | HWRWRYAAAYA |
| *Foxa_known3* | - | 8.8 | 16.7 | VMWRWGTMAAYA |
| *Foxc1_1* | - | 1.7 | 13.6 | NBNVHGTAAAYAAASR |
| *Foxd1_1* | - | 3.8 | 15.8 | SYWRWGTAAACADWVS |
| *Foxd1_2* | - | 1.9 | 13.9 | GTAAACAW |
| *Foxf1* | - | -0.1 | 11.9 | YRHAYAAACAHNB |
| *Foxj1_1* | - | -1.2 | 10.7 | TAAACAAACAHWD |
| *Foxj1_2* | - | 8.4 | 16.2 | HNNRTAAACAMWNNND |
| *Foxk1* | - | 7.6 | 17.8 | NDWWKRTAAACAAWVNB |
| *Foxo_1* | - | 1.0 | 13.0 | VWMAACAMNHH |
| *Foxo_2* | - | 4.2 | 16.2 | MWBGTAAACAAMVN |
| *Foxo_3* | - | 3.3 | 15.3 | WRWAAAYAAVWD |
| *Foxo_4* | - | 2.2 | 14.0 | KGWAAACA |
| *HDAC2_disc2* | + | 1.7 | 13.3 | WRRGYMAAYA |
| *Pou5f1_disc1* | - | 9.9 | 10.8 | YWTTSWYATGCARAW |
| *Pou5f1_known1* | + | 9.1 | 10.6 | YWTTSWYATGCWRAT |
| *Sox_6* | + | 3.9 | 10.2 | WAWSWMM |
| *TCF12_disc2* | + | 7.4 | 11.9 | WVWGYMMAYA |
| *p300_disc3* | - | 3.7 | 14.3 | VWRDRYAMAYA |

**Table S4:** List of transcription factors affected by rs356182. List based on the HOCOMOCO database, filtered for strong interactions only.

| | **SCORE** | | **P-VALUE** | | |
|---|---|---|---|---|---|
| **GENE** | **REF** | **ALT** | **REF** | **ALT** | **SEQUENCE MATCH** |
| *FOXA1* | 5.103 | 6.044 | 0.00078 | 4.29E-06 | TTGCAATGTAAACAAATCTTC |
| *FOXA2* | 5.138 | 6.065 | 0.00053 | 7.15E-07 | TTTGCAATGTAAACAAATCTTCT |
| *FOXA3* | 4.240 | 5.158 | 0.00289 | 8.70E-06 | CTTTGCAATGTAAACAAATCTTCTC |
| *FOXC1* | 6.499 | 7.356 | 0.00023 | 6.60E-06 | TGCTTTGCAATGTAAACAAATCTTCTCCA |
| *FOXD1* | 5.239 | 6.088 | 0.00046 | 5.57E-06 | TGCTTTGCAATGTAAACAAATCTTCTCCA |
| *FOXF2* | 5.383 | 6.098 | 0.00012 | 1.67E-06 | TTGCAATGTAAACAAATCTTC |

| | | | | | |
|---|---|---|---|---|---|
| *FOXO1* | 5.248 | 6.180 | 0.00109 | 8.47E-06 | TGCTTTGCAATGTAAACAAATCTTCTCCAG |
| *FOXO3* | 4.976 | 5.913 | 0.00096 | 3.13E-07 | CTTTGCAATGTAAACAAATCTTCTC |
| *FOXO4* | 4.628 | 5.560 | 0.00202 | 4.20E-05 | GCAATGTAAACAAATCT |
| *FOXP2* | 4.380 | 5.247 | 0.00127 | 7.63E-06 | GCAATGTAAACAAATCT |

**Table S5:** Gene filtering statistics. Adjusted P-value < 0.05; absolute fold-change > 2. LUHMES gene universe, n=24,774, 13,918 mapped to GO terms. Simplification by combining redundant terms with >70% overlap between genes.

| *Condition* | *Gene expression direction change* | *Total genes in set (FC>2)* | *Genes mapped to GO terms* | *Terms enriched for mapped genes- full* | *Simplified term list* |
|---|---|---|---|---|---|
| A/- vs A/G | + | 801 | 510 | 287 | 119 |
| | - | 752 | 464 | 203 | 96 |
| G/- vs A/G | + | 730 | 558 | 577 | 210 |
| | - | 639 | 324 | 33 | 22 |

**Table S6:** Top 25 GO terms associated with the changed genes in each condition. Simplified ontology to reduce redundant terms.

***A/- vs A/G; up-regulated gene annotations***

| *ID* | *Description* | *Gene-Ratio* | *Bg-Ratio* | *q-value* |
|---|---|---|---|---|
| GO:0051962 | positive regulation of nervous system development | 53/510 | 481/13918 | 2.07e-09 |
| GO:0045666 | positive regulation of neuron differentiation | 43/510 | 345/13918 | 3.03e-09 |
| GO:0050808 | synapse organization | 44/510 | 380/13918 | 1.38e-08 |
| GO:0007409 | axonogenesis | 47/510 | 437/13918 | 2.14e-08 |
| GO:0050804 | modulation of chemical synaptic transmission | 44/510 | 392/13918 | 2.14e-08 |
| GO:0099177 | regulation of trans-synaptic signaling | 44/510 | 393/13918 | 2.14e-08 |
| GO:0050769 | positive regulation of neurogenesis | 46/510 | 428/13918 | 2.61e-08 |
| GO:0061564 | axon development | 49/510 | 477/13918 | 2.61e-08 |
| GO:0010720 | positive regulation of cell development | 48/510 | 487/13918 | 1.53e-07 |
| GO:0007416 | synapse assembly | 25/510 | 157/13918 | 1.89e-07 |
| GO:0007610 | behavior | 48/510 | 498/13918 | 2.65e-07 |
| GO:0048167 | regulation of synaptic plasticity | 25/510 | 165/13918 | 4.61e-07 |
| GO:0050807 | regulation of synapse organization | 27/510 | 203/13918 | 1.76e-06 |
| GO:0050803 | regulation of synapse structure or activity | 27/510 | 213/13918 | 4.64e-06 |
| GO:0007411 | axon guidance | 30/510 | 257/13918 | 4.8e-06 |
| GO:0097485 | neuron projection guidance | 30/510 | 258/13918 | 4.92e-06 |
| GO:0042391 | regulation of membrane potential | 36/510 | 364/13918 | 1.28e-05 |
| GO:0098742 | cell-cell adhesion via plasma-membrane adhesion molecules | 27/510 | 227/13918 | 1.39e-05 |
| GO:0034329 | cell junction assembly | 36/510 | 371/13918 | 1.85e-05 |
| GO:0006935 | chemotaxis | 42/510 | 475/13918 | 2.06e-05 |
| GO:0042330 | taxis | 42/510 | 477/13918 | 2.2e-05 |
| GO:0034765 | regulation of ion transmembrane transport | 37/510 | 395/13918 | 2.61e-05 |
| GO:0051963 | regulation of synapse assembly | 16/510 | 91/13918 | 2.69e-05 |
| GO:0034762 | regulation of transmembrane transport | 41/510 | 466/13918 | 2.8e-05 |

| | | | | |
|---|---|---|---|---|
| GO:0023061 | signal release | 39/510 | 440/13918 | 4.31e-05 |

***A/- vs A/G; down-regulated gene annotations***

| *ID* | *Description* | *Gene-Ratio* | *Bg-Ratio* | *q-value* |
|---|---|---|---|---|
| GO:0048598 | embryonic morphogenesis | 41/464 | 491/13918 | 0.000199 |
| GO:0048568 | embryonic organ development | 32/464 | 363/13918 | 0.000869 |
| GO:0008015 | blood circulation | 35/464 | 425/13918 | 0.000869 |
| GO:0001525 | angiogenesis | 34/464 | 411/13918 | 0.000869 |
| GO:0060688 | regulation of morphogenesis of a branching structure | 10/464 | 45/13918 | 0.000912 |
| GO:0007423 | sensory organ development | 37/464 | 479/13918 | 0.000912 |
| GO:0019932 | second-messenger-mediated signaling | 29/464 | 330/13918 | 0.000912 |
| GO:0003013 | circulatory system process | 35/464 | 444/13918 | 0.000912 |
| GO:0043114 | regulation of vascular permeability | 9/464 | 38/13918 | 0.00122 |
| GO:0042330 | taxis | 36/464 | 477/13918 | 0.00135 |
| GO:0007610 | behavior | 37/464 | 498/13918 | 0.00135 |
| GO:0070830 | bicellular tight junction assembly | 11/464 | 64/13918 | 0.00191 |
| GO:0048017 | inositol lipid-mediated signaling | 18/464 | 163/13918 | 0.00191 |
| GO:0009791 | post-embryonic development | 12/464 | 77/13918 | 0.00191 |
| GO:0003007 | heart morphogenesis | 21/464 | 213/13918 | 0.00191 |
| GO:0006935 | chemotaxis | 35/464 | 475/13918 | 0.00191 |
| GO:0030198 | extracellular matrix organization | 26/464 | 303/13918 | 0.00191 |
| GO:0120192 | tight junction assembly | 11/464 | 66/13918 | 0.00191 |
| GO:0043062 | extracellular structure organization | 26/464 | 304/13918 | 0.00191 |
| GO:0007204 | positive regulation of cytosolic calcium ion concentration | 21/464 | 216/13918 | 0.00191 |
| GO:1901342 | regulation of vasculature development | 24/464 | 271/13918 | 0.00213 |
| GO:0002009 | morphogenesis of an epithelium | 34/464 | 463/13918 | 0.00213 |
| GO:0120193 | tight junction organization | 11/464 | 69/13918 | 0.00223 |
| GO:0048514 | blood vessel morphogenesis | 35/464 | 487/13918 | 0.00223 |
| GO:0050866 | negative regulation of cell activation | 16/464 | 140/13918 | 0.00223 |

***G/- vs A/G; up-regulated gene annotations***

| *ID* | *Description* | *Gene-Ratio* | *Bg-Ratio* | *q-value* |
|---|---|---|---|---|
| GO:0043588 | skin development | 33/558 | 213/13918 | 7.49e-08 |
| GO:0030198 | extracellular matrix organization | 38/558 | 303/13918 | 5.13e-07 |
| GO:0043062 | extracellular structure organization | 38/558 | 304/13918 | 5.13e-07 |
| GO:0072091 | regulation of stem cell proliferation | 16/558 | 58/13918 | 5.3e-07 |
| GO:0048568 | embryonic organ development | 41/558 | 363/13918 | 1.25e-06 |
| GO:0008544 | epidermis development | 33/558 | 253/13918 | 1.25e-06 |
| GO:0140014 | mitotic nuclear division | 34/558 | 270/13918 | 1.54e-06 |
| GO:0050673 | epithelial cell proliferation | 37/558 | 318/13918 | 2.42e-06 |
| GO:0048608 | reproductive structure development | 39/558 | 351/13918 | 3.04e-06 |
| GO:0061458 | reproductive system development | 39/558 | 355/13918 | 3.76e-06 |
| GO:1905818 | regulation of chromosome separation | 15/558 | 63/13918 | 6.05e-06 |
| GO:0000819 | sister chromatid segregation | 26/558 | 184/13918 | 6.22e-06 |
| GO:0045787 | positive regulation of cell cycle | 37/558 | 336/13918 | 6.3e-06 |
| GO:0051302 | regulation of cell division | 22/558 | 137/13918 | 6.3e-06 |
| GO:0090068 | positive regulation of cell cycle process | 31/558 | 254/13918 | 7.56e-06 |
| GO:0044784 | metaphase/anaphase transition of cell cycle | 14/558 | 58/13918 | 1.03e-05 |
| GO:0010965 | regulation of mitotic sister chromatid separation | 14/558 | 59/13918 | 1.12e-05 |
| GO:0044057 | regulation of system process | 46/558 | 488/13918 | 1.12e-05 |
| GO:0050678 | regulation of epithelial cell proliferation | 32/558 | 275/13918 | 1.12e-05 |
| GO:0000070 | mitotic sister chromatid segregation | 23/558 | 157/13918 | 1.24e-05 |
| GO:0070371 | ERK1 and ERK2 cascade | 28/558 | 223/13918 | 1.37e-05 |

| | | | | |
|---|---|---|---|---|
| GO:0051983 | regulation of chromosome segregation | 18/558 | 101/13918 | 1.43e-05 |
| GO:0007423 | sensory organ development | 45/558 | 479/13918 | 1.43e-05 |
| GO:0033045 | regulation of sister chromatid segregation | 16/558 | 81/13918 | 1.58e-05 |
| GO:0051306 | mitotic sister chromatid separation | 14/558 | 62/13918 | 1.63e-05 |

***G/- vs A/G; down-regulated gene annotations***

| *ID* | *Description* | *Gene-Ratio* | *Bg-Ratio* | *q-value* |
|---|---|---|---|---|
| GO:0007156 | homophilic cell adhesion via plasma membrane adhesion molecules | 20/324 | 150/13918 | 8.16e-07 |
| GO:0007600 | sensory perception | 33/324 | 427/13918 | 2.05e-06 |
| GO:0007606 | sensory perception of chemical stimulus | 14/324 | 81/13918 | 4.42e-06 |
| GO:0098742 | cell-cell adhesion via plasma-membrane adhesion molecules | 22/324 | 227/13918 | 1.17e-05 |
| GO:0007608 | sensory perception of smell | 10/324 | 43/13918 | 2.26e-05 |
| GO:0050804 | modulation of chemical synaptic transmission | 28/324 | 392/13918 | 6.14e-05 |
| GO:0099177 | regulation of trans-synaptic signaling | 28/324 | 393/13918 | 6.14e-05 |
| GO:0042391 | regulation of membrane potential | 26/324 | 364/13918 | 0.000145 |
| GO:0007411 | axon guidance | 21/324 | 257/13918 | 0.000193 |
| GO:0097485 | neuron projection guidance | 21/324 | 258/13918 | 0.000193 |
| GO:0061564 | axon development | 29/324 | 477/13918 | 0.000615 |
| GO:0008038 | neuron recognition | 8/324 | 44/13918 | 0.00161 |
| GO:0008037 | cell recognition | 12/324 | 108/13918 | 0.00172 |
| GO:0007409 | axonogenesis | 26/324 | 437/13918 | 0.0024 |
| GO:0050906 | detection of stimulus involved in sensory perception | 11/324 | 97/13918 | 0.00275 |
| GO:0007610 | behavior | 28/324 | 498/13918 | 0.00275 |
| GO:0006935 | chemotaxis | 27/324 | 475/13918 | 0.00304 |
| GO:0042330 | taxis | 27/324 | 477/13918 | 0.00309 |
| GO:0051606 | detection of stimulus | 17/324 | 228/13918 | 0.00375 |
| GO:0007214 | gamma-aminobutyric acid signaling pathway | 5/324 | 17/13918 | 0.00466 |
| GO:0009593 | detection of chemical stimulus | 9/324 | 72/13918 | 0.00573 |
| GO:0071805 | potassium ion transmembrane transport | 14/324 | 171/13918 | 0.0061 |
| GO:0021559 | trigeminal nerve development | 4/324 | 11/13918 | 0.0104 |
| GO:0001508 | action potential | 11/324 | 117/13918 | 0.0106 |
| GO:0016339 | calcium-dependent cell-cell adhesion via plasma membrane cell adhesion molecules | 6/324 | 35/13918 | 0.0155 |

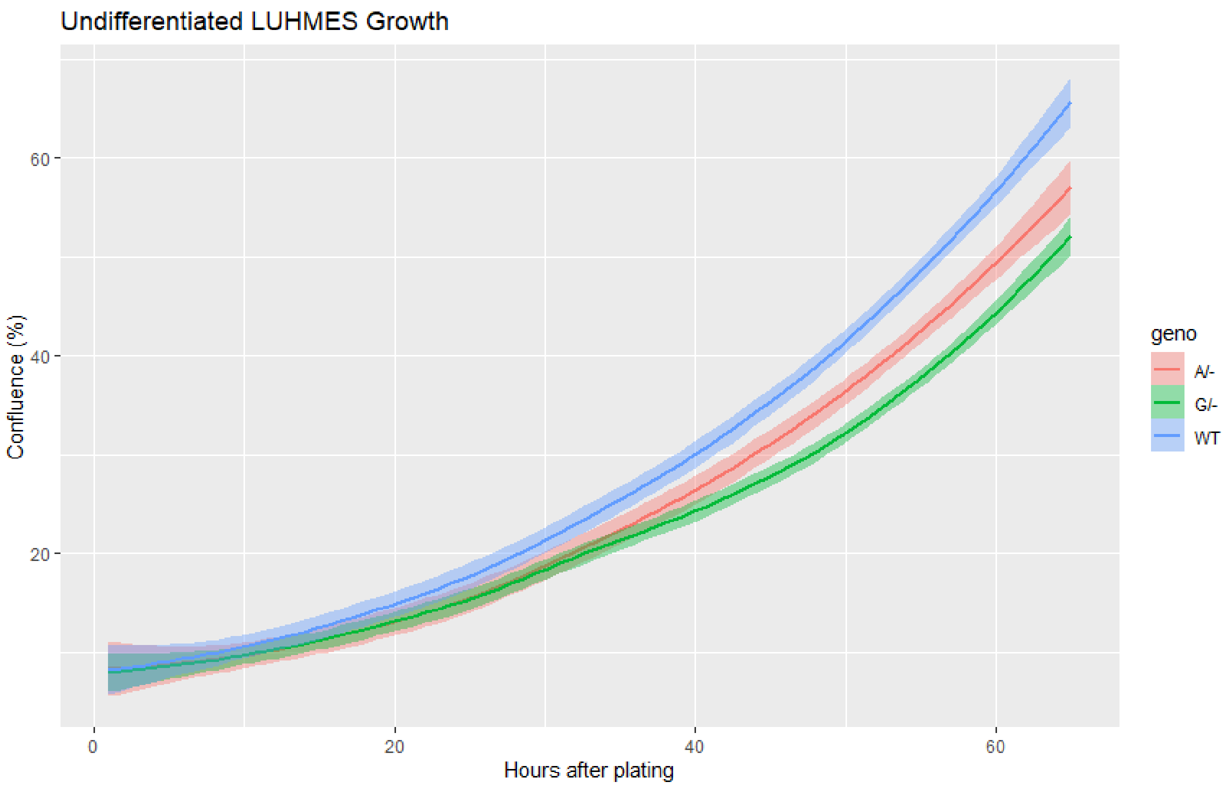


**Figure S2:** Incucyte growth rate data of lesion clones. Time after plating on the X-axis and percent confluence on the Y-axis.

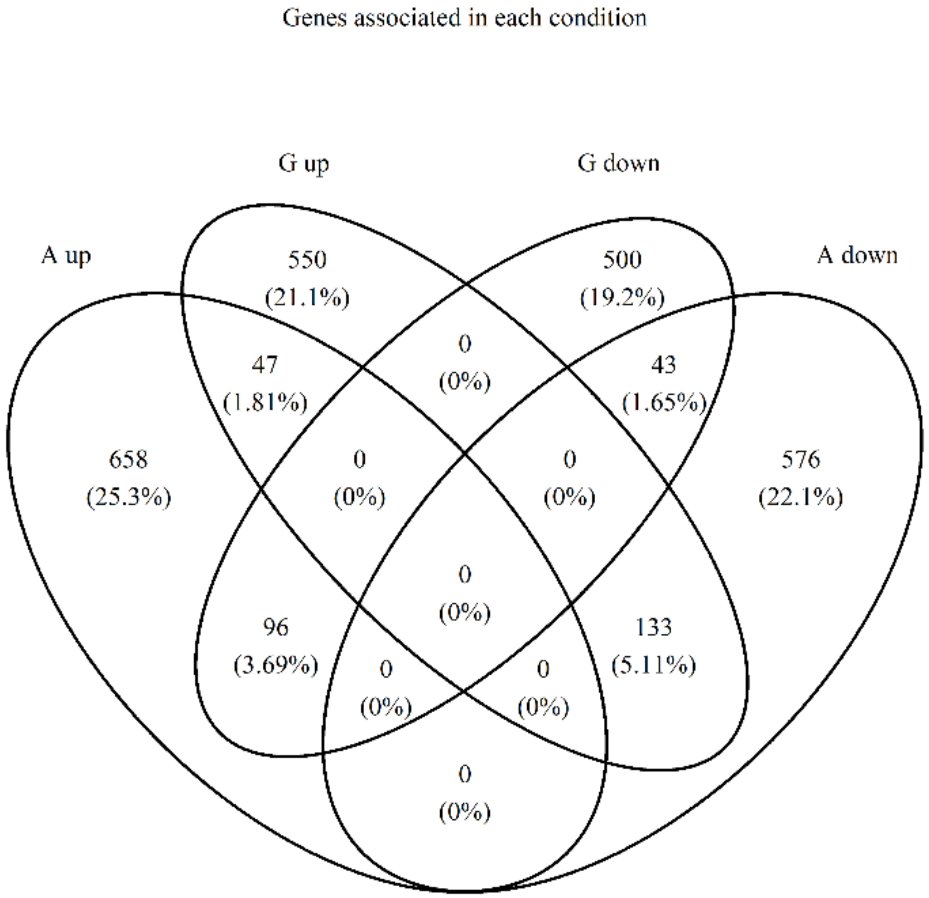


**Figure S3**: Venn diagram showing the unique and overlapping genes in each condition.
